# Eco-evolutionary Dynamics of Colonization: When Does Source Selection Matter?

**DOI:** 10.64898/2026.10.29.762850

**Authors:** Eve Nancy Rowland, Helen J. Wearing, Davorka Gulisija

## Abstract

Ancestral balancing selection may facilitate evolutionary rescue following migration into a new habitat by providing founder populations with adaptive alleles, even when those alleles were not the fittest in the ancestral habitat. However, recent studies emphasize the roles of immigration history and admixture on colonization success. The extent to which balanced polymorphism may contribute to colonization success and the conditions under which it matters remain unclear. We use computer simulations to investigate how the mode of selection in the source population influences colonization success in new habitats under eco-evolutionary population dynamics, different invasion scenarios, and different selection regimes in the new habitat. We model source populations under balancing selection, directional selection, and neutrality, and allow founder groups of different sizes to invade a novel stable or perturbed habitat. We find that balancing selection in ancestral populations leads to higher rates of establishment in new habitats than are observed under source neutrality or directional selection when founder populations are relatively small, which would occur with a population bottleneck. This trend is observed regardless of whether invaders experience stable or variable conditions in a new habitat and persists under increasingly harsh conditions or greater environmental perturbations. When the colonizing populations are large, selection in the source population has little effect on invasion success. This study elucidates the interaction between genetic diversity and propagule pressure on colonization success, highlighting the number of adaptive alleles as a unique metric.

## INTRODUCTION

Habitat destruction, environmental shifts, and human-mediated transport are increasingly introducing organisms into new habitats (le Roux and McGeoch 2008; Hufbauer et al. 2011). Extreme environmental shifts, such as the severe weather events resulting from climate change, are expected to further push populations to their adaptive limits (Thuiller 2004; Holt et al. 2005; Thomas 2010; Usinowicz and Levine 2021) and increase the number of habitat invaders (Hufbauer et al. 2011). Yet not all migrants successfully colonize new habitats (Capellini et al. 2015; Estoup et al. 2016; Kołodziejczyk et al. 2025). Some studies have hypothesized that elevated genetic diversity in migrants facilitates invasion success (Barrett and Schluter 2008; Lee and Gelembiuk 2008; Forsman 2014; Posavi et al. 2014; Promy et al. 2023), while others emphasize the key role of introduction history or admixture (Roman and Darling 2007; Blackburn et al. 2015; Capellini et al. 2015; Jaspers et al. 2021; Kołodziejczyk et al. 2025).

However, the roles of these mechanisms have not been jointly evaluated, and the relative importance of genetic diversity for colonization success remains unclear. Here, we use an eco-evolutionary computational model to examine how genetic diversity levels in a source population affect the probability of establishment during habitat colonization across a range of invasion scenarios and novel selection regimes.

Colonization success refers to immigration, establishment, and persistence in a new habitat (MacArthur and Wilson 1967). It is often attributed to introduction history, particularly propagule pressure, which refers to the cumulative effect of founding population size and number of colonization events (Roman and Darling 2007; Blackburn et al. 2015; Jaspers et al. 2021). Populations with high propagule pressure, either from larger founding population sizes and/or multiple introduction events, are less vulnerable to demographic stochasticity, which increases colonization success (Blackburn et al. 2015). Additionally, large populations are more likely to carry or give rise to diverse alleles (Blackburn et al. 2015), which may provide adaptive alleles that may be necessary for a colonization event. Repeated introductions essentially produce the same effect by increasing the founding population size. Additionally, founder populations can increase their likelihood of establishment by increasing their genetic diversity via admixture from independent populations (Roman and Darling 2007). For populations with relatively low genetic diversity, admixture among divergent populations in the invasive range may be a key contributor to establishment success (Kołodziejczyk et al. 2025).

There are limitations to the impact that population size can have on invading populations. Propagule pressure alone does not explain all cases of establishment. Some invasions succeed despite bottlenecks (Kołodziejczyk et al. 2025), whereas others fail despite repeated introductions. This suggests that colonization success may reflect interactions among introduction history, ecological release, phenotypic plasticity, and adaptive evolution (Lee 2002; Hovick and Whitney 2019; Kołodziejczyk et al. 2025). The benefit of admixture is also limited and can facilitate adaptation only when beneficial alleles are already present in the contributing populations. It is likely that a wide range of factors, including de novo mutations in novel habitats, founding population sizes, number of colonization events, and preexisting genetic diversity, all influence the success of establishment in novel habitats.

Timely adaptation to a new stable habitat or a new regime of temporal variation may be pivotal for population persistence. It has long been hypothesized that populations harboring pre-existing elevated genetic diversity have an advantage when experiencing novel conditions and are more likely to establish in novel habitats (Carroll and Dingle 1996; Gomulkiewicz and Holt 1995; Barrett and Schluter 2008; Lee and Gelembiuk 2008; Posavi et al. 2014). This can be particularly beneficial during rapid environmental change, since adaptation can occur faster from existing variation than from de novo mutations (Barrett and Schluter 2008; Matuszewski et al. 2015; Guzella et al. 2018; Gulisija 2026). Standing genetic variation can arise from various forms of balancing selection, including heterozygote advantage, direct negative frequency-dependent selection, and selection that varies across time and/or space (Hedrick 2007).

Interestingly, it has also been proposed that invaders are more likely to originate from habitats characterized by spatial or temporal environmental heterogeneity (Lozon and MacIsaac 1997; Lee and Bell 1999; Lee and Gelembiuk 2008), further pointing to the role of varying environments. It follows that temporally varying balancing selection in the source population may predispose migrants to succeed in colonization. Although this possibility has long been proposed, balancing selection was not considered likely.

Balancing selection has been considered to occur in a limited set of scenarios, such as overdominance or direct negative frequency dependence (Felsenstein 1976; Gulisija 2026). In temporally varying environments, overdominance arises in geometric mean over time, conditions for which are considered generally weak (Gulisija 2026). However, recent theoretical work has identified a broader class of conditions under which balancing polymorphism can arise in temporally varying environments, through different models of storage effect (Ellner and Hairston Jr. 1994; Gulisija and Kim 2015; Svardal 2015; Gulisija et al. 2016; Yamamichi and Hoso 2017). The storage effect, initially proposed in community ecology (Chesson and Warner 1981; Chesson 1983; Warner and Chesson 1985), promotes genetic diversity by ‘storing’ alleles in a portion of the population that experiences partial protection from varying selection, such as in a particular life stage (e.g., seed banks) (Ellner and Hairston Jr. 1994; Kirkpatrick and Peischl 2013) or a subpopulation in a different habitat patch (Gulisija and Kim, 2015). In addition, the storage effect increases population persistence under eco-evolutionary models in habitats experiencing temporal variation, leading to evolutionary rescue (Rowland and Gulisija 2025).

Evolutionary rescue in the new habitat refers to a phenomenon where populations persist due to adaptation to new conditions, thus allowing the population to survive conditions where it would perish otherwise (Bell and Gonzalez 2009; Bell 2013; Barfield and Holt 2016; Xu et al. 2023). Genetic variation can promote evolutionary rescue (Martin et al. 2013; Schiffers et al. 2013) by providing the material needed for rapid adaptation (Barrett and Schluter 2008; Lee and Remfert 2007; Rowland and Gulisija 2025). Facilitating evolutionary rescue is important during invasions since populations may be forced to adapt rapidly within the novel habitat to conditions that vary significantly from their founding population conditions (Lee and Remfert 2007). For example, polymorphism-facilitated evolutionary rescue is important for vulnerable populations, such as those at the expanding edge of a species’ range (Pease et al. 1989; Bowler and Benton 2005; Hetem et al. 2014; Holt et al. 2005, 2022). Although evolutionary rescue theory examines how adaptation can promote population persistence, relatively little work has explicitly considered how evolutionary rescue may contribute to colonization success in novel habitats.

The role of source-population diversity in colonization success has not been theoretically evaluated alongside invasion history and novel selection regimes. In particular, it remains unclear when elevated standing variation promotes establishment, how it interacts with founding population size, and whether diversity maintained by balancing selection is differently beneficial in stable versus temporally varying novel environments. We use an in silico forward-in-time eco-evolutionary population framework to examine how the source population’s diversity affects establishment in a novel habitat under different invasion scenarios and new selection regimes.

We model range expansions from populations with elevated, neutral, and low levels of genetic diversity. Elevated-diversity populations are characterized by balanced polymorphism arising under the storage effect in the ancestral habitat, whereas low-diversity populations are drawn from habitats in which source populations experienced directional selection in a direction other than in the new habitat. We evaluate establishment under the two novel selection regimes: directional selection or temporally varying selection, and across a range of founding population sizes. This framework allows us to examine how source-population evolutionary history shapes colonization success and how its effects interact with founding population size.

## MODEL AND METHODS

We model eco-evolutionary dynamics in a founding population that migrates to a new habitat. The founders are haploid, characterized by a single biallelic locus, with alleles *d* and *b*, originating from the source population evolving under population dynamics and a distinct selection regime from that of the new habitat. To represent different immigration scenarios, we vary the number of migrants originating from the source population, *N*_founder_, eco-evolving under (i) directional selection favoring the *d* allele, (ii) temporally varying balancing selection due to the spatial storage effect, where *b* and *d* are quasi-neutral (Hartl and Cook 1973), or (iii) genetic drift (see *Supplementary Methods* for details on how ancestral genotypes were generated using the model developed by Rowland and Gulisija (2025)) (*Figure S1*). In the new habitat, the alleles are deleterious, *d*, and beneficial, *b*, in their effect on fitness.

We simulate the eco-evolutionary dynamics of a founder population in a new habitat forward in time (Kim and Weihe 2009). During each discrete generation, a founder population experiences eco-evolution due to the effects of recurrent mutation at rate μ, new environmental conditions generating either directional or temporally varying selection, and stochastic reproduction following the logistic growth function. For clarity, *Table 1* defines the symbols used in the study.

**Table 1.**
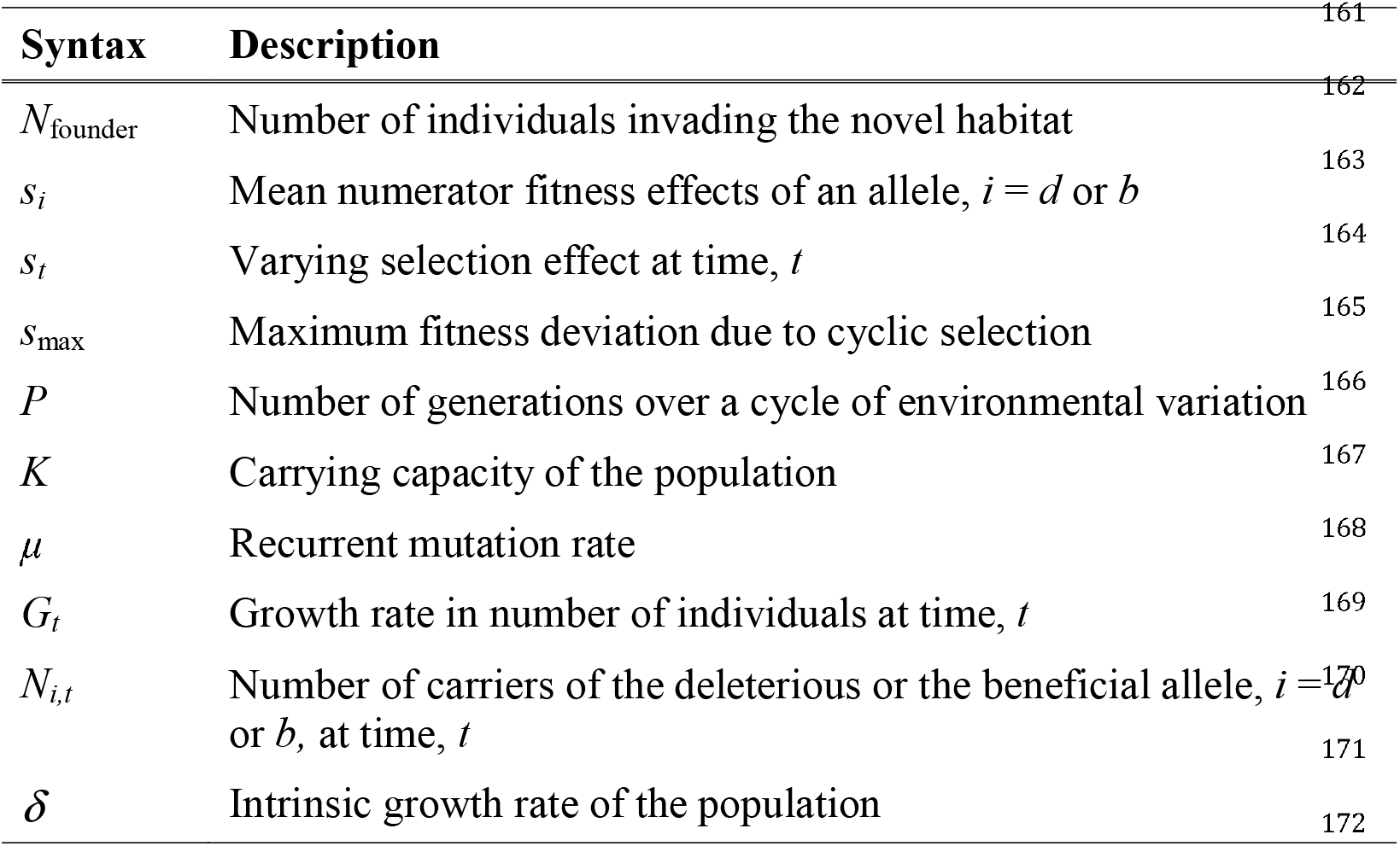
Mathematical Symbols.

The effects of the new selective environment are implemented through genotype-based fitness effects, while population dynamics is modeled via carrying-capacity-dependent growth rates, *G*_*t*_. Then, the number of individuals carrying the deleterious allele (*d*) and the number of individuals carrying the beneficial allele (*b*) after density-regulated selection are determined by:

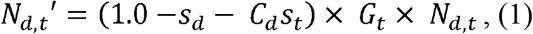

And

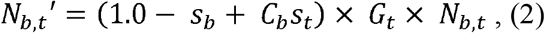

where *s*_*t*_ is the varying selection effect, *s*_*i*_ is the mean numerator fitness effects of an allele (*i* = *d* or *b*) in the new habitat, and *N*_*d,t*_ and *N*_*b,t*_ are the numbers of carriers of *d* and *b* allele before density-regulated selection. *C*_*d*_ and *C*_*b*_ are constants that modify the magnitude of fitness oscillations, while *G*_*t*_ represents the population growth rate.

In a stable new habitat, under directional selection, *s*_*t*_ = 0. When the selection in a new habitat varies over time:

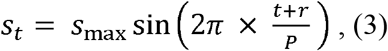

here *s*_max_ is the maximum departure from mean fitness over the cycle of fitness oscillations and *P* represents the number of generations occurring within the cycle of variation. We first consider *C*_*d*_ *= C*_*b*_ = 1, where both alleles experience fitness oscillations of equal magnitude. We then extend the model to unequal magnitudes by allowing either *C*_*d*_ ≠ 1 or *C*_*b*_ ≠ 1, while keeping the other fixed at one.

The growth rate of the population at time *t*:

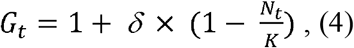

where *δ* is the intrinsic growth rate, and *K* is the carrying capacity of the population, with *N*_*t*_, = *N*_*d,t*_ + *N*_*b,t*_, is the population size at time *t*.

In order to simulate an increasingly perturbed novel habitat over time, the varying selection coefficient increases incrementally over time, reaching the maximum at 1.5*s*_max_:

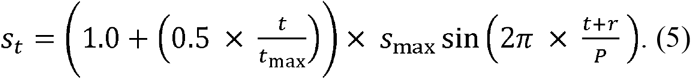

Finally, the number of offspring in the next generation for each genotype is sampled from a Poisson distribution with means of *N*_*d,t*_ and *N*_*b,t*_.

### Implementation

We eco-evolved 20,000 replicate populations for each parameter combination (ancestral-invasion scenario) initiated from a founder group of size *N*_founder_ and at a carrying capacity *K*, for a maximum time of 100 times the carrying capacity (100*K*) generations. We simulated a factorial design in which each population first evolved under one of the three ancestral conditions: directional selection, balancing selection, or drift; after which *N*_founder_ migrants were introduced into a novel habitat and evolved under one of two new selection regimes: stable directional selection or temporally varying selection around a new optimum (*Figure 1*).

**Figure 1.**
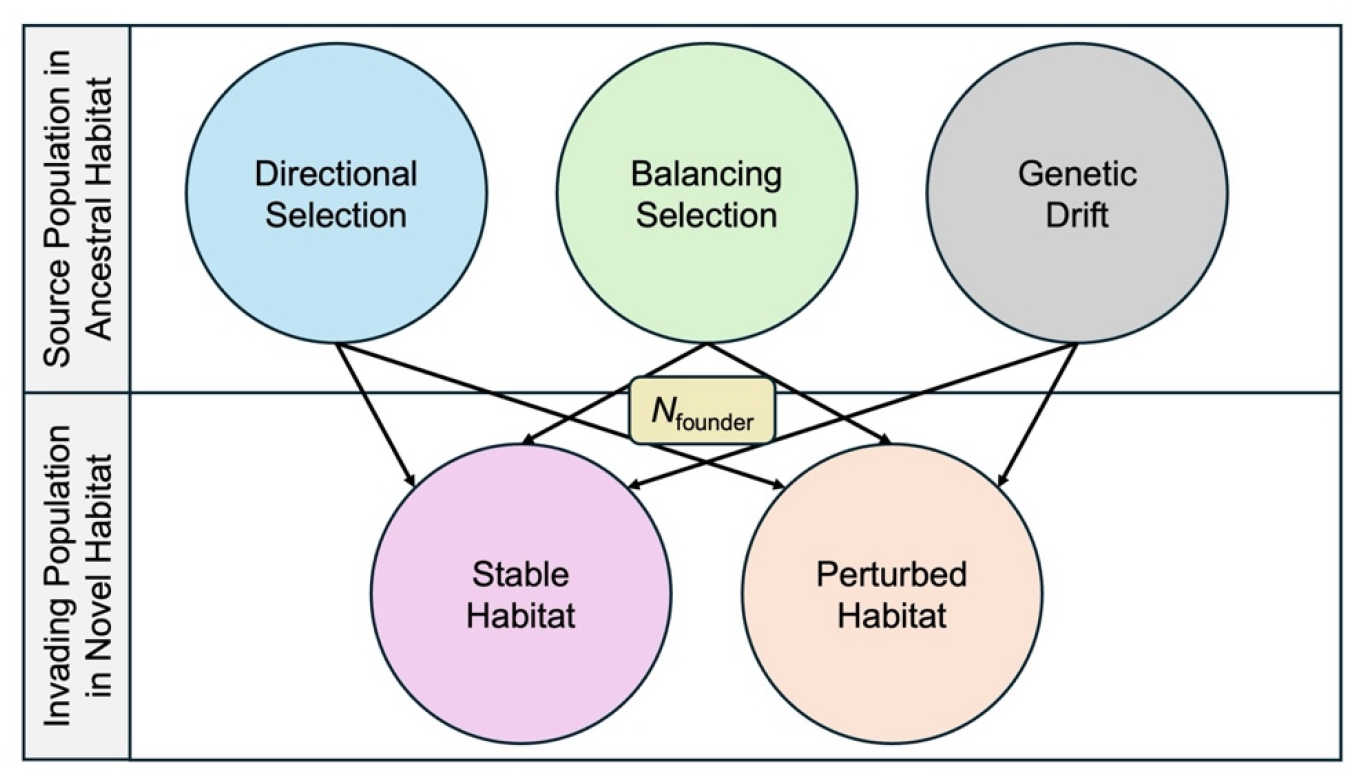
Implementation figure. The founders originated from ancestral populations evolving under (i) directional selection against *b* allele (*h*_model_/*h*_drift_ < 1.0), (ii) balancing selection due to the storage effect (*h*_model_/*h*_drift_ > 1.5), and (iii) genetic drift (*h*_model_/*h*_drift_ = 1.0) as a control, with *h*_model_/*h*_drift_ being the level of observed heterozygosity compared to that under drift. Each founding population of size *N*_founder_ migrates into one of two novel environments: (i) a stable habitat under directional selection favoring the *b* allele or (ii) a perturbed habitat where temporally varying selection occurs around a new mean favoring *b*.

To ensure the appropriate initial distribution of ancestral levels of diversity, the frequencies of the two alleles in founder populations were sampled from the frequency distributions generated in source populations which either exhibited low diversity due to directional selection against *b* (*s* = -0.01), elevated diversity from temporally varying balancing selection resulting in elevated heterozygosity (*P* = 12; *P* = 24; *P* = 48; all with, *s*_max_ = 0.2), or genetic drift, as a control. The expected levels of heterozygosity compared to that under drift, *h*_model_/*h*_drift_, in the balanced source populations were 1.90 for *P* = 12, 3.02 for *P* = 24, 4.78 for *P* = 48, where *P* indicates the period of temporally varying environments in ancestral habitat (see *Supplemental Methods*), but 0.51 for those that arose in stable habitats under directional selection.

Stable conditions in the novel habitat were modeled over a range of allelic mean fitnesses with *s*_*d*_ ranging from -0.55 to -0.8 and *s*_*b*_ ranging from -0.15 to -0.4, both in increments of -0.01. The invasion of a perturbed habitat was modeled with *s*_*d*_ = -0.55, *s*_*b*_ = -0.15 and *s*_max_ = 0.2, and with an increasing magnitude of fitness oscillations following equation [5]. Founder population sizes ranged from *N*_founder_ = 100 to 4000 at increments of 50 individuals.

Extinction was considered as the population size falling below 25 individuals, 1/4 of the smallest *N*_founder_. The extinction threshold is the population size below which populations can no longer recover and eventually go extinct. A lower threshold allows higher establishment success because it gives populations more time to adapt, and vice versa (*Figure S2*). However, establishment success declines more rapidly as the threshold increases for founders originating from low-diversity conditions than for those from high-diversity conditions (see *Figure S2* for an assessment of the robustness of the results to this metric). This indicates that the relatively low threshold that was chosen to generate the results of this study reveals general principles and, if anything, represents a conservative scenario.

The carrying capacity (*K*) for the majority of invasion scenarios was 4,000 individuals, while we examined the robustness of the observed trends to lower carrying capacities under *K* = 2000, 1000, and 500 individuals (*Figure S6*). Recurrent mutation occurred at a rate of μ*K* = 0.05 or μ = 0.0000125 per generation. Populations that established in a new habitat were those that survived to the maximum simulation time of 100*K* generations. Adaptation in a new habitat is considered to occur by a quasi-fixation of the beneficial allele, *b* (frequency > 0.99). The intrinsic growth rate of populations, *δ* was 0.5.

## RESULTS

The number of migrants and the level of ancestral heterozygosity interactively shape invasive success in novel habitats. When migrant numbers are low, the level of genetic diversity is critical for establishment; conversely, when genetic diversity is low, migrant number becomes critical. However, when either genetic diversity or migrant number is sufficiently high, populations establish successfully regardless of the other factor (*Figure 2*). This pattern holds whether invasions occur in stable or perturbed habitats (*Figures 3 & 4*), across extinction thresholds (*Figure S2*), and even with increased harshness against the non-adaptive genotype (*d, Figure 5*), or both genotypes (*Figure S3*), thus potentially causing rapid post-invasion decline. The interaction also persists in novel habitats where the magnitude of temporally varying selection increases over time (*Figure S4*), or where two alleles experience different magnitudes of fitness oscillations (*Figure S5*). Finally, the results are robust to assumptions about the carrying capacity (*Figure S6*).

**Figure 2.**
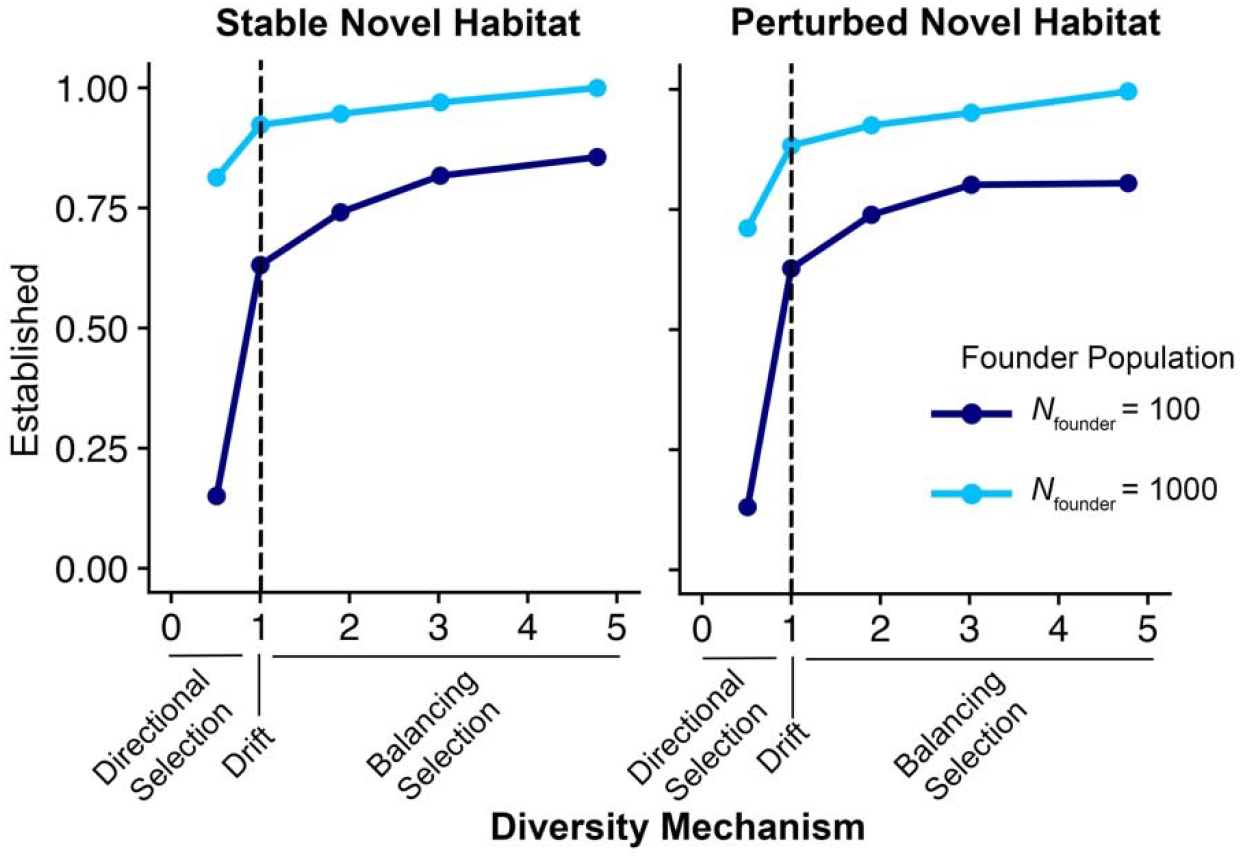
Relationship between ancestral genetic diversity and establishment in a novel habitat. Successful establishment requires higher levels of ancestral genetic diversity in smaller founder populations (*N*_founder_ = 100, dark blue) than in larger founder populations (*N*_founder_ = 1000, light blue). This pattern holds regardless of whether the novel selection regime is stable (left) or perturbed (right). Ancestral genetic diversity was simulated under conditions of balancing selection via the spatial storage effect (*h*_model_*/h*_drift_ = 1.90 for *P* = 12, *h*_model_*/h*_drift_ = 3.02 for *P* = 24, *h*_model_*/h*_drift_ = 4.78 for *P* = 48), directional selection (*h*_model_*/h*_drift_ = 0.51), or drift (*h*_model_*/h*_drift_ = 1.0). The populations within the novel habitats experienced either directional selection (stable novel habitat on left; *s*_*d*_ = -0.55, *s*_*b*_ = -0.15 with *s*_max_ = 0.0) or a perturbed novel habitat with temporally varying selection (perturbed habitat on right; *s*_*d*_ = -0.55, *s*_*b*_ = -0.15 with *s*_max_ = 0.2). Simulations were conducted across 20,000 replicate runs with carrying capacity *K* = 4,000 over 400,000 generations. Recurrent mutation occurred at a rate of μ = 0.0000125 per generation.

**Figure 3.**
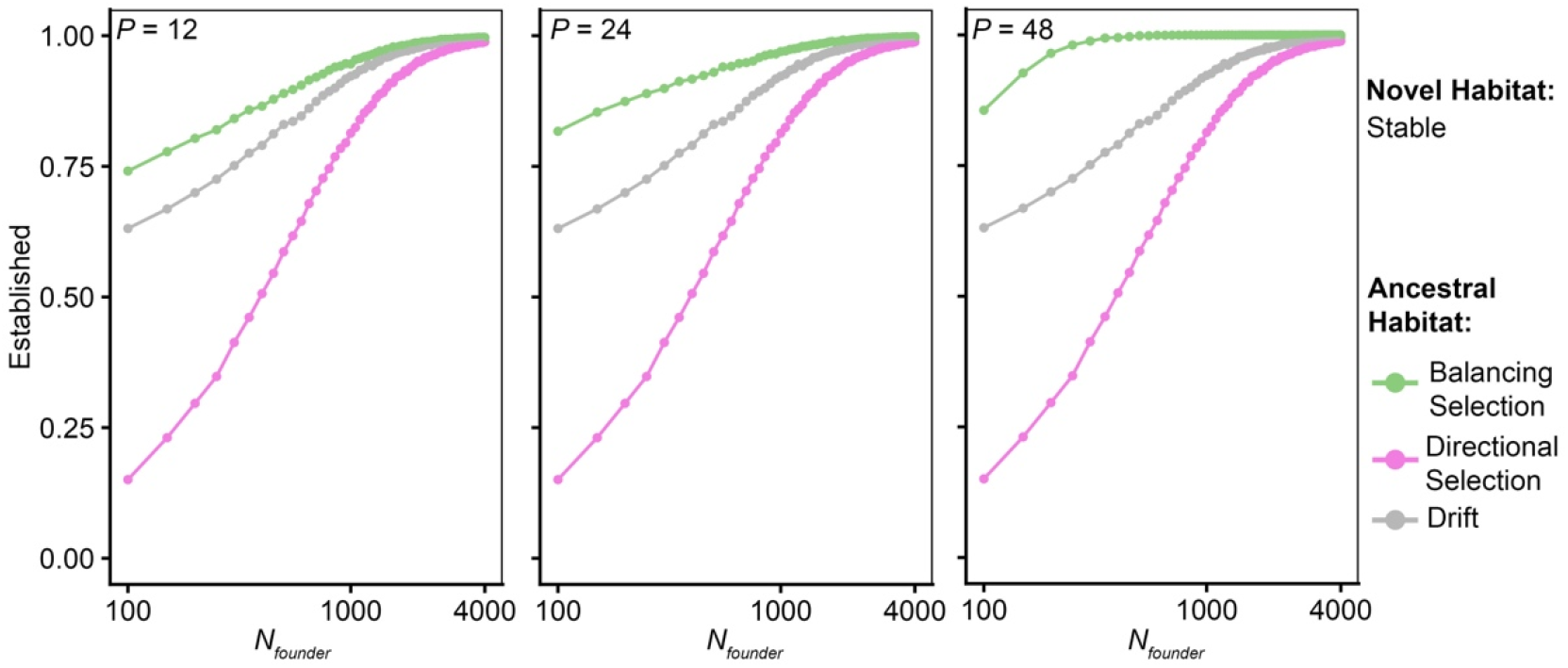
Probability of establishment in a stable novel habitat as a function of founder population size, *N*_founder_. Founder populations originated under conditions of balancing selection via the spatial storage effect (light green, *h*_model_*/h*_drift_ = 1.90 for *P* = 12, *h*_model_*/h*_drift_ = 3.02 for *P* = 24, *h*_model_*/h*_drift_ = 4.78 for *P* = 48), directional selection favoring *d* allele (pink, *h*_model_*/h*_drift_ = 0.51), or drift (light grey), and experienced stable conditions favoring *b* allele (*s*_*d*_ = -0.55, *s*_*b*_ = -0.15 with *s*_max_ = 0.0) in the novel habitat. The *N*_founder_ = 100 to 4000 at increments of 50 individuals (left to right). Simulations were conducted over 20,000 replicate runs with *K* = 4,000 over 400,000 generations. Recurrent mutation occurs at a rate such that μ = 0.0000125 per generation. The x-axis is shown on a log scale.

**Figure 4.**
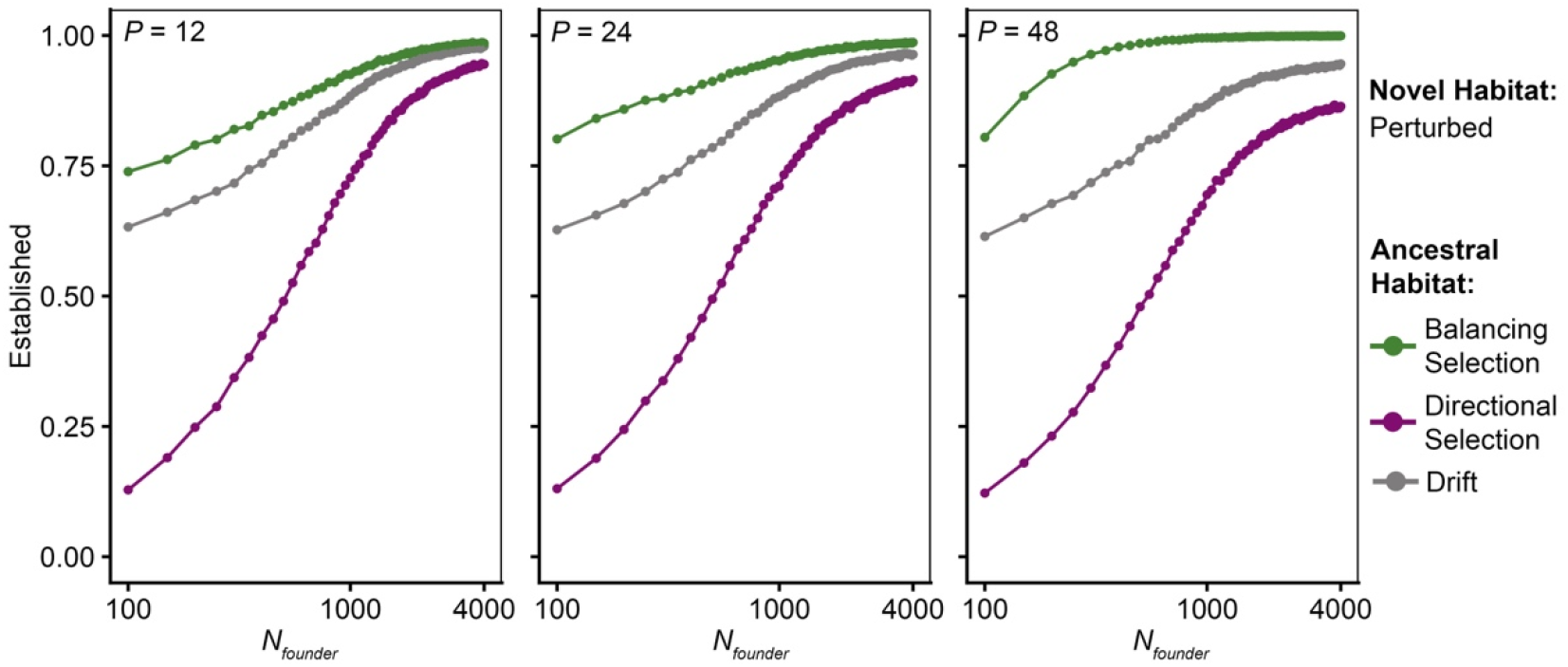
Probability of establishment in a perturbed novel habitat under temporally varying selection as a function of the founder population size, *N*_founder_. Founder populations originated under conditions of balancing selection under the spatial storage effect (dark green, *h*_model_*/h*_drift_ = 1.90 for *P* = 12, *h*_model_*/h*_drift_ = 3.02 for *P* = 24, *h*_model_*/h*_drift_ = 4.78 for *P* = 48), directional selection favoring *d* (purple, *h*_model_*/h*_drift_ = 0.51), or drift (dark grey), and experienced a perturbed novel habitat around new optima favoring *b* (*s*_*d*_ = -0.55/*s*_*b*_ = -0.15 with *s*_max_ = 0.2) in the novel habitat. Simulations were conducted over 20,000 replicate runs with *K* = 4,000 over 400,000 generations. The *N*_founder_ = 100 to 4000 at increments of 50 individuals (left to right). Recurrent mutation occurs at a rate such that μ = 0.0000125 per generation. The x-axis is shown on a log scale.

Taken together, these results suggest that populations originating from those that experienced balancing selection have a higher probability of colonizing novel habitats through evolutionary rescue following bottlenecks than those originating under directional purifying selection or neutrality.

### Invasions into stable habitats

To understand how ancestral selection impacts invasion success, we allowed founder groups across a range of sizes (*N*_founder_ = 100 to 4000) to migrate to a novel stable habitat where selection favors the *b* allele, which was not favored in the ancestral habitat. Across a wide range of founding population sizes, populations that originate from habitats under the balancing selection establish at a higher rate than otherwise (*Figure 3*). The founding populations arising from populations with the highest level of heterozygosity (*P* = 48) resulted in the highest rate of establishment; over 95% of the replicate populations reached establishment with a founding population size of only 200 individuals or higher. The groups with lower ancestral heterozygosity typically required somewhat larger sizes to reach the same levels of establishment. Particularly, at founder-population sizes as small as 100 individuals, the populations originating from ancestral balanced habitats had a higher probability of successful invasion compared to those originating under directional selection against *b* (i.e. purifying selection under stable ancestral conditions; ∼80% probability compared to 15%). This increased probability of successful establishment demonstrates that genetic diversity within the ancestral population can allow otherwise more vulnerable invaders, which undergo bottlenecks, an increased probability of successful invasion. This suggests that these immigrants benefited from carrying a larger number of adaptive genotypes, allowing them to escape extinction under conditions in which establishment would otherwise fail (*Figure S7*).

### Invasions into perturbed habitats

The type of selection in the new habitat had little effect on the patterns described above. The founder populations (*N*_founder_ = 100 to 4000) that originated from a habitat under the storage effect were more likely to successfully establish within a perturbed novel habitat, where fitness oscillates around a new optimum favoring *b*, than those arising from populations under purifying selection, or drift (*Figure 4*). This benefit of ancestral balancing selection is, again, especially apparent at lower population sizes. When a founder population is initiated at 100 individuals, the probability of establishment in the cases of ancestral balanced populations is ∼80% as opposed to only ∼13% in populations coming from stable habitats. Likewise, the populations with the highest level of genetic variation in the ancestral population (*P* = 48) had an establishment probability of above 95% for founder populations above 300 individuals, while the highest probability ever reached for populations arising from stable habitats was 86% at *N*_founder_ = 4000. Thus, ancestral balancing selection broadly provides opportunities for populations to undergo evolutionary rescue in new habitats.

### Robustness to model perturbations

The trends described above clearly illustrate the interactions between diversity level and the number of migrants in shaping invasion success, indicating that balancing selection might play a key role in colonization following bottlenecks. Here, we demonstrate that these patterns are robust across a wide range of model assumptions.

To examine whether the effects of environmental harshness depend on ancestral selection history, we expanded our model beyond the baseline scenario of migration from one environmental optimum to another. Specifically, we increased the fitness disadvantage to the deleterious allele, *d*, by examining the dynamics under a range of selection coefficients *s*_*d*_, from -0.55 to -0.8 by increments of -0.01. This allowed us to examine increasingly strong selection against *d*, which removes the maladapted portion of the founder population more rapidly.

Despite this stronger selection pressure, the overall trends remained consistent: founder diversity continued to play an important role in establishment success and tended to retain more stable levels of establishment compared to the purifying selection and drift (*Figure 5*). In contrast, the populations originating from the stable habitat with directional selection against the *b* allele showed the fastest decline in establishment as selection against *d* increased. These results indicate that the advantage of ancestral balancing selection persists even under harsher environmental conditions. In fact, as environments became more hostile, the differences between founders originating from balanced versus stable ancestral conditions became more pronounced (*Figure 5*).

**Figure 5.**
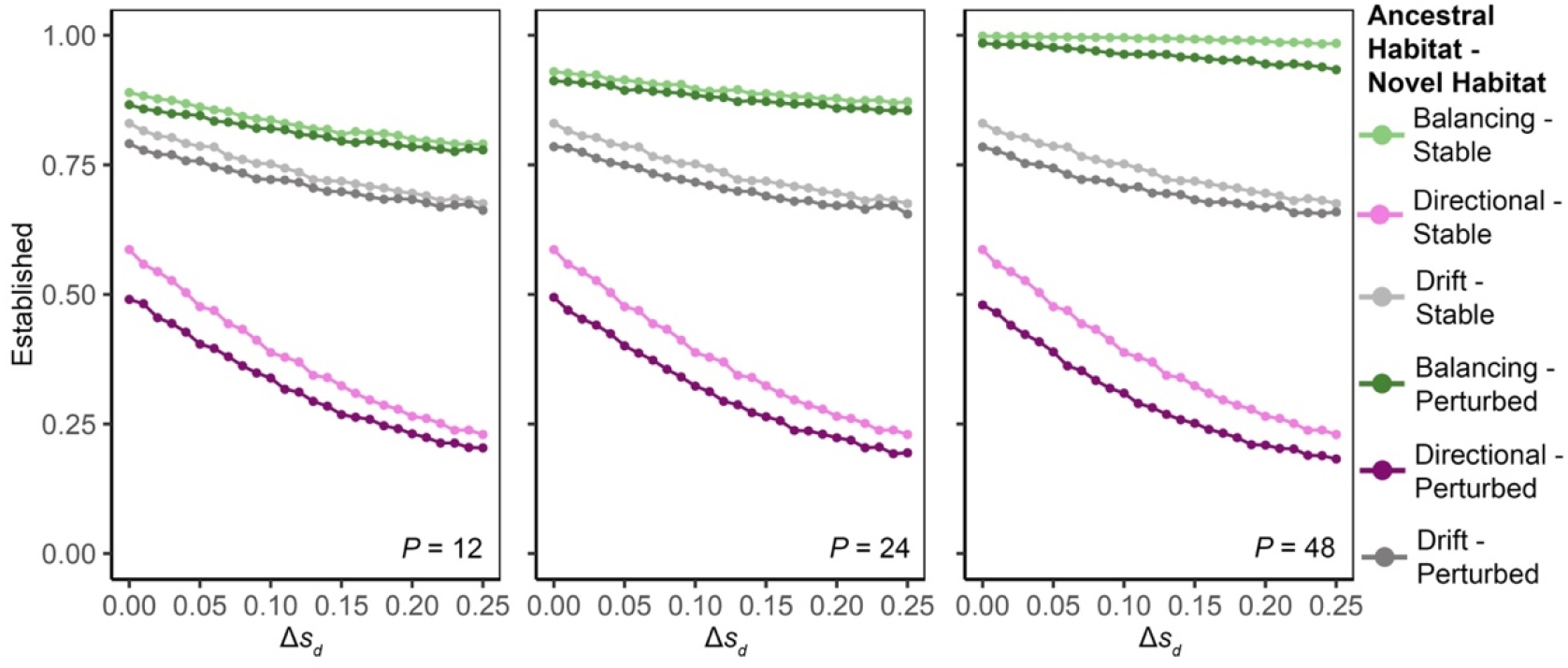
Probability of establishment in a novel habitat with stronger selection against the *d* allele. Simulations were conducted over 20,000 replicate runs with *K* = 4,000 and a founder population size, *N*_founder_ = 500. Recurrent mutation occurs at a rate of μ = 0.0000125 per generation. Founder populations originated under conditions of balancing selection under the spatial storage effect (light green/dark green, *h*_model_*/h*_drift_ = 1.90 for *P* = 12, *h*_model_*/h*_drift_ = 3.02 for *P* = 24, *h*_model_*/h*_drift_ = 4.78 for *P* = 48), directional selection against *b* allele (pink/purple, *h*_model_*/h*_drift_ = 0.51), or drift (light grey/dark grey). The novel environment was either stable (light green/pink/light grey) with directional selection (*s*_*d*_ = -0.55, *s*_*b*_ = -0.15 with *s*_max_ = 0.0) or perturbed (dark green/purple/dark grey) with temporally varying selection (*s*_*d*_ = -0.55, *s*_*b*_ = -0.15 with *s*_max_ = 0.2), both favoring the *b* allele. To test different levels of environmental harshness, we set *s*_*d*_ to -0.55 to -0.8, in increments of 0.01, the x-axis shows the corresponding change in selection, Δs_d_, with increasing Δs_d_ indicating greater environmental harshness from left to right.

To reduce the buffering effect of the beneficial subpopulation while simultaneously generating harsher conditions through selection against *d*, we next decreased the fitness of both alleles, with *s*_*d*_ ranging from -0.55 to -0.8 and *s*_*b*_ ranging from -0.15 to -0.4 by increments of -0.01. The ancestral balancing selection continued to increase colonization success, even as extinction became more likely (*Figure S3*).

Habitats that populations migrate into may experience shifts over time. To test whether balancing selection in founding populations also provides an advantage in a novel habitat that changes over time and becomes increasingly perturbed, we increased the magnitude of fitness oscillations over the course of the simulation. The founding populations originating from conditions favoring the storage effect still had the highest rate of establishment compared to the other populations (*Figure S4*). The populations with the highest starting level of heterozygosity (*P* = 48) had the highest rate of establishment across all conditions, but even the populations with lower levels of starting heterozygosity (*P* = 12 and 24) had a higher rate of establishment than the populations under drift and those under purifying selection in the ancestral habitat.

Another difference that colonizers may experience could be unequal fitness oscillations between the two alleles. Across a range of unequal magnitudes of fitness oscillations between the two alleles, the balanced founding populations maintained a higher rate of establishment than those under directional selection or drift (*Figure S5*).

The trends above were robust to the assumptions about the carrying capacity (*Figure S6*), as long as colonization was possible. Overall, these results provide strong support for the role of ancestral balancing selection during colonization bottlenecks. The interaction between diversity level and the number of migrants clearly points to the importance of the number of adaptive genotypes that appear in the novel habitat.

### Intuition

An interactive relationship between the ancestral diversity level, which is a function of the frequency of the beneficial genotype (*p*.), and the total founder population size (*N*.) is expected, since *N*_*b,t*_ = *p*_*t*_*N*_*t*_, the number of adaptive genotypes in the founder population. As founders enter a new habitat, the subpopulation of maladapted genotypes (*d*) is expected to decline while the subpopulation of adapted genotypes (*b*) increases. The key to colonization success is for *N*_*b,t*_, to reach a critical size before *N*_*t*_ declines below the extinction threshold. Then, a large founding population, even with low *p*_*t*_, may have sufficient time for selection to build *N*_*b,t*_ to the critical size, while a small group founded through a bottleneck depends on the number of adaptive genotypes already being present.

Consider the expected change in the size of the subpopulations of carriers for each genotype, *d* and *b*, in a new stable environment under the eco-evolutionary population dynamics given in equations [1] and [2], *N*^*′*^_*b,t*_ = (1 − *s*_*d*_) × *G*_*t*_ × *N*_*d,t*_ and *N*^*′*^_*b,t*_ = (1 − *s*_*b*_) × *G*_*t*_ × *N*_*b,t*_. Here, *s*_*i*_ is the mean numerator fitness effect of an allele (*i* = *d* or *b*), a genotype-based relative fitness deviation at time *t* due to selection, *s*_*d*_ ∈ (0,1) is the selective cost to the maladapted genotype, *s*_*d*_ > *s*_*b*_ > 0 gives the selective advantage to the adapted genotype, and *G*_*t*_ is the density-dependent population growth rate. Total size is *N*_*t*_ = *N*_*d,t*_ + *N*_*b,t*_, and the frequency of the fitter type is 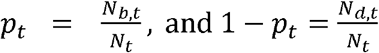. Then,

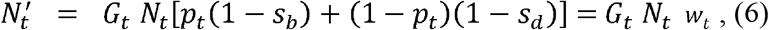

with *w*_*t*_ the mean population fitness at time *t*.

The population grows in the next generation if

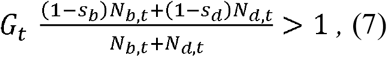

when

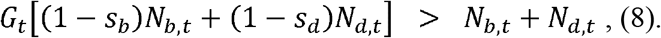

Thus, the new contribution to the next generation made by the fitter type *G*_*t*_ (1 − *s*_*d*_) *N*_*b,t*_ must exceed the declining numbers due to *G*_*t*_ (1 − *s*_*d*_) *N*_*d,t*_. The dynamics at the threshold of extinction, thus, depend on a critical number of adaptive genotypes, *N*_*b,t*_, i.e. the product of the frequency of the beneficial allele, *p*_*t*,_ and population size *N*_*t*_.

In this model, a shift in the abundances of two genotypes leads to evolutionary rescue.

While the *N*_*d*_ subpopulation declines, the *N*_*b*_counterbalances this effect by an increase, thereby stabilizing the overall population size and enabling populations to persist at a higher rate than would be possible in the absence of polymorphism. We illustrate this with a series of stochastic invasions into a novel habitat (*Figure S7*). When colonists arrive in numbers already near the extinction threshold, those that evolved in a stable environment where selection acted against the adaptive allele *b* carry too few adaptive individuals (pink). As a result, the population typically cannot rise above the threshold before the maladapted group (green) declines, and extinctions are frequent. In contrast, colonists drawn from a balanced ancestral population usually include enough adaptive individuals to trigger evolutionary rescue, letting the total population grow instead of collapsing.

## DISCUSSION

The long-standing debate over whether colonization success is driven primarily by genetic diversity or by propagule pressure overlooks the key point of their interaction: the absolute number of adaptive genotypes that arrive in the novel habitat. This is particularly important for smaller founding populations that rely almost exclusively on the adaptive variation they already possess compared to larger founding populations that may survive long enough for new beneficial mutations to emerge and increase in frequency. The population establishes by reaching a critical mass of adaptive alleles before demographic decline becomes irreversible, either via evolution from low adaptive diversity or de novo mutation (less common in small populations), or because immigrants already carry adaptive alleles that can sweep in the new habitat.

Therefore, experiencing balancing selection in an ancestral population, such as the storage effect, greatly increases the probability that successful adaptation and establishment will occur when founder populations are small. Our study, thus, demonstrates that the conditions within an ancestral population, specifically the level of ancestral diversity, can have a profound impact on colonization success.

Populations may respond to environmental change through adaptation or dispersal to avoid extinction (Pease et al.1989; Bowler and Benton 2005; Holt et al. 2005; Duputié et al. 2012; Hetem et al. 2014; Holt et al. 2022). In some populations, dispersal will increase the chance of survival compared to adaptation in the existing environment (Souza et al. 2023). In fact, non-indigenous species on a global scale are undergoing range expansion at an increasing rate as a direct result of climate change (Rius et al. 2014). Previous studies have suggested that invaders may arise from habitats that vary temporally and/or spatially (Lee and Bell 1999).

Additionally, seasonal fluctuations (similar to those modeled in this study) likely impact the rate at which a population is able to adapt to a novel habitat (Lee and Bell 1999). Other studies have reported a strong association between disturbance and colonization (Lozon and MacIsaac 1997), suggesting that invaders are more likely to come from perturbed habitats. In particular, landscape fragmentation is becoming more common as anthropogenic climate change leads to the alteration of habitats (Boeye et al. 2013; Mona et al. 2014), which will probably lead to increased migration out of these fragmented habitats. Our study clarifies the circumstances under which balancing selection in ancestral habitats promotes evolutionary rescue in novel habitats. This benefit appears particularly important for small founding population sizes, such as following bottlenecks. We further clarify that the size of the founder population is the key factor in determining the effects of ancestral conditions on invasive success.

Admixture between genetically divergent populations is common among successful invaders (Kołodziejczyk et al. 2025). This points to genetic diversity and rapid adaptation as prerequisites for colonization. Yet admixture alone cannot pinpoint the original source of adaptive alleles, nor can it reveal how much ancestral balancing selection contributed to that variation. Addressing these uncertainties requires genomic surveys of the source populations that produced both successful and failed invasions. Yet, genetic diversity is typically measured only in already-established invaders (Kołodziejczyk et al. 2025), where it may have been reshaped by the invasion process itself. During rapid adaptation, genome-wide variation can rise if many loci respond to selection (Hayward and Sella 2022) or shrink as adaptive alleles become fixed.

Elevated diversity in an invader may therefore be a consequence of establishment rather than a reflection of its ancestral state. Conversely, low genome-wide diversity after a founder event can coexist with a “critical mass” of adaptive alleles at key loci, a pattern that may fuel the genetic paradox of invasions (Allendorf and Lundquist 2003). Our theoretical results suggest that even severely bottlenecked founders can succeed if they carry balanced polymorphisms for traits under selection, making the absolute number of adaptive genotypes at arrival, not overall heterozygosity, the key variable. Empirical evidence supports this view: both propagule size and standing variation improve establishment odds (Hovick and Whitney 2019).

Indeed, colonization success is likely the result of an interplay of several factors acting in concert. For example, phenotypic plasticity adds a further layer to the debate. Because plastic genotypes can express different phenotypes when environments shift (Pigliucci et al. 2006; Schrieber and Lachmuth 2017), plasticity can, like a large founder population, provide crucial time for genetic adaptation (Lande 2009). Plastic responses tend to evolve in disturbed habitats and can also be enhanced by admixture between divergent populations (Gulisija and Newberry 2026). A full understanding of evolutionary rescue in novel environments must therefore integrate founder size, the ancestral pool of adaptive alleles, and phenotypic plasticity, which was not featured in our study. Time-series genomic sampling of source and colonist populations would be especially powerful for disentangling their respective roles.

We modeled ancestral balance under the spatial storage effect, which requires migration and spatial heterogeneity in varying selection between demes. Another form of the storage effect, the genomic storage effect, can arise in uniform varying environments and unstructured populations due to genetic recombination and phenotypic plasticity (Gulisija et al. 2016), which is likely to arise in perturbed habitats. Incorporating genetic plasticity into future models will clarify how genetic diversity and plasticity together affect colonization success.

Another consideration is that we modeled eco-evolutionary dynamics at a focal locus directly involved in adaptation and evolutionary rescue. We did not consider genome-wide fitness effects, which may influence population persistence in the long run, independently of adaptation to the immediate environmental change. In particular, population bottlenecks can reduce genetic diversity across the genome and contribute to population decline even when adaptation occurs at the focal locus. Such effects may help explain the prominent role that admixture often plays in successful colonization by restoring genome-wide genetic variation that was lost during bottlenecks.

Our study, however, focused on the immediate dynamics of evolutionary rescue at a locus critical for adaptation and suggests that, at least during the initial stages of colonization, the number of adaptive alleles available to selection may be a key determinant of persistence. More comprehensive computational studies that jointly consider polygenic fitness across all contributing traits, demographic bottlenecks, balancing selection at key loci, admixture, and phenotypic plasticity will be needed to determine how interactions among these processes ultimately shape the probability of successful range expansion.

## Acknowledgments

We would like to thank the members of the Gulisija Computational Genetics and Evolution Lab for their assistance throughout this study. We would like to thank Dr. Matthew J. Campen, Dr. Joseph A. Cook, and Dr. Marcus A. Garcia for their thoughtful input on the manuscript. This work was supported by startup funds from the University of New Mexico and by the National Science Foundation DEB-2622346 to DG.

## Data availability

The code used to generate the results will be made available on GitHub upon manuscript publication.

## Author Contributions

DG and ENR conceived the study and designed the model and simulation code. ENR generated the data and figures. All authors interpreted the findings and co-wrote the article.

## Statements and Declarations

The authors declare no conflict of interest.

