## Supplemental Material for "Eco-evolutionary Dynamics of Colonization: When Does Source Selection Matter?"

**Supplemental Information**

***Supplementary Methods***

To simulate ancestral conditions, we used the computational model and code from Rowland and Gulisija (2025). Within the ancestral habitat, the storage effect was established in a subdivided population across two demes. The first deme evolved under neutrality and acted as a refuge (R) from the negative effects of selection, while the other deme experienced temporally varying selection (V) determined by a sine function with a selective advantage or disadvantage at time *t* based on the fitness effect of each allele (*s_t_*) and a maximum fitness deviation in the ancestral population of *s*_max_, as described in equation [3] in Model and Methods.Each generation, the demes undergo stochastic mutation under recurrent rate *µ*, stochastic gene flow, and deterministic changes in genotype sizes, followed by stochastic reproduction.

The genotype sizes change due to the varying selection (*s_t_*), the mean environmental effect (environmental harshness, *S_i_*) and the deme’s growth rate (*G_i,t_*), such that:

$N_{id,t}'=\left( 1-S_{i}- s_{t} \right)\times G_{i,t}\times N_{id,t}$,

and

$N_{ib,t}'=\left( 1- S_{i}+ s_{t} \right)\times G_{i,t}\times N_{ib,t}$ ,

where *N_id,t_* and *N_ib,t_* represent the number of individuals carrying the *d* allele and the *b* allele in deme *i =* R or V. When a deme is under drift, *s_t_* = 0 and for the deme under directional selection in the ancestral habitat, *s_t_* = -0.01. The migrant frequencies are sampled from the perturbed habitat, i.e. deme V. Population size change is based on the logistic growth method, as described in Model and Methods. The intrinsic growth rate was 0.5 for all demes. Reproduction occurs stochastically. The number of individuals with each of the two alleles in each population in the next generation is sampled from a Poisson distribution with mean *N_id,t_* and mean *N_ib,t_*.

In order to detect the storage effect, the level of heterozygosity was measured. The expected heterozygosity in each deme at time *t* is:

$h_{i,t}=2 p_{i,t}q_{i,t}$,

where *p_i,t_* is the frequency of the *d* allele, and *q_i,t_* is the frequency of the *b* allele, in a deme *i* at time *t*. The weighted heterozygosity at time *t* across V and R is, thus,

$h_{t}=\frac{2\times p_{V,t}q_{V,t}\times N_{V,t}}{N_{V,t}+ N_{R,t}}+ \frac{2\times p_{R,t}q_{R,t} \times N_{R,t}}{N_{V,t}+ N_{R,t}}$.

We report expected heterozygosity relative to neutral expectation under identical mutation and migration rates, such that:

$h (model/drift)=\frac{\Sigma_{n}\frac{{\Sigma_{t}h}_{model, t}}{T_{\mathrm{model}}}/n}{\Sigma_{n}\frac{{\Sigma_{t}h}_{drift, t}}{T_{\mathrm{drift}}}/n}$,

where *n* is the number of replicate simulations and *T.* represents expected survival time under temporally varying selection or drift. Three parameter combinations were selected to be used for this study: *P* = 12 with *M* = 250; *P* = 24 with *M* = 100; *P* = 48 with *M* = 75; all with, *s*_max_ = 0.2, providing a range of levels of diversity. The level of heterozygosity compared to that under drift, *h*_model_/*h*_drift_, was 1.90 for *P* = 12, 3.02 for *P* = 24, 4.78 for *P* = 48. *M* represents the number of individuals that are exchanged between the demes at carrying capacity.

For all simulations in this study, the mean environmental effect was *S*_V_ = 0.35 and *S*_R_ = 0. The rate of recurrent mutation was *μ* = 0.0000125. The rate of recurrent mutation was *μ* = 0.0000125. The carrying capacity (*K*) for both R and V was 4,000 individuals, and the maximum simulation time is 400,000 generations across 8,000 replicates.

***Supplemental Figures***

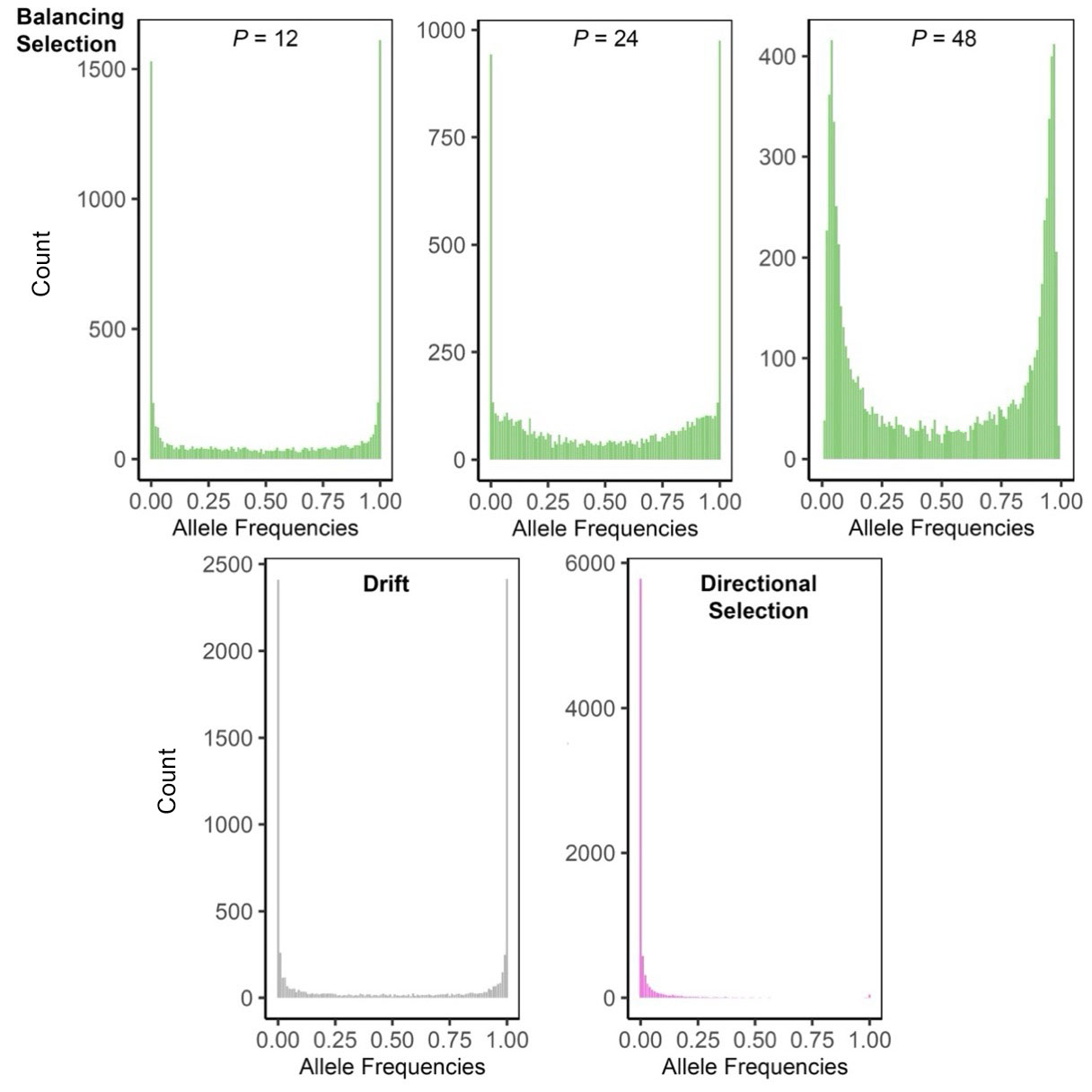

**Figure S1. Equilibrium distributions of the beneficial (*b*) allele frequencies at the onset of invasion under different ancestral selection regimes.** The distributions were generated assuming a recurrent mutation rate *μ* = 0.0000125 or *μK* = 0.05 per generation, a carrying capacity of 4,000 for each of the two populations that exchange migrants, and by conducting 8,000 replicate simulation runs with a total time of 400,000 generations. At the onset of each simulation, the starting allele frequencies in founders in the novel habitat were assigned by sampling from these distributions of equilibrial frequencies in deme V. The distribution where directional selection acted against the ancestral population was generated using the selection pressure of -0.01 against the *b* allele. The distributions where balancing selection via the storage effect was present in the ancestral population assumed *s*_max_ = 0.2 in one subpopulation and drift in the other, and migration rates of *M* = 250 for *P* = 12, *M* = 100 for *P* = 24, and *M* = 75 for *P* = 48.

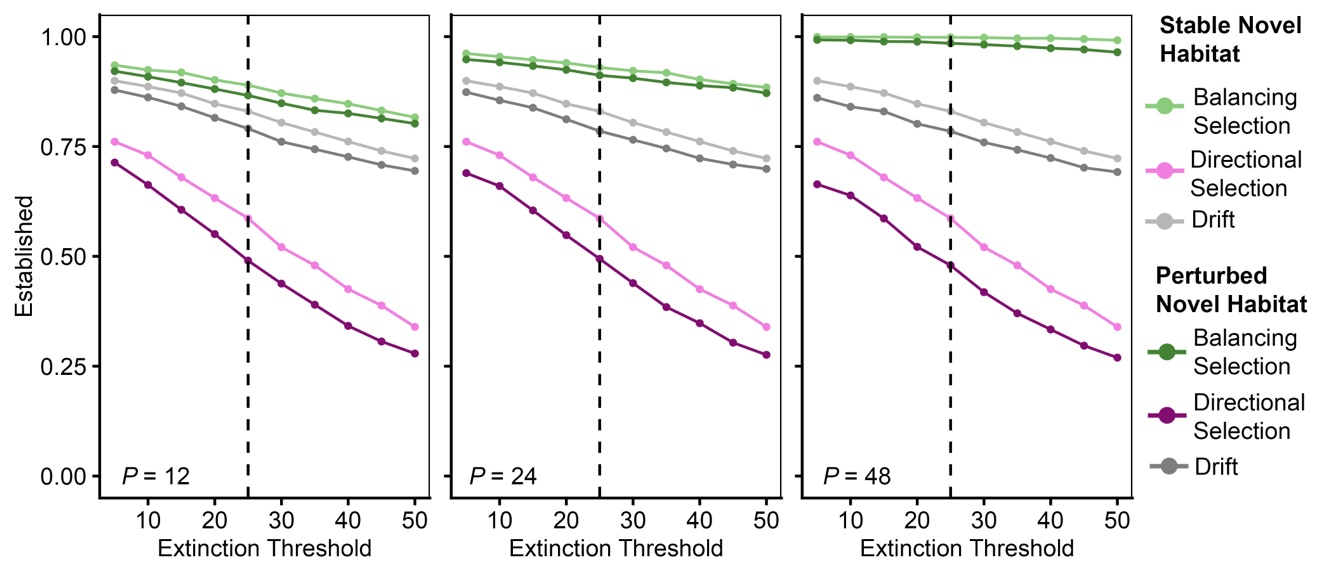

**Figure S2. The probability of establishment in a novel habitat under differing extinction thresholds**. The extinction thresholds tested were *N* = 5 – 50 individuals (in increments of 5). The dashed line marks *N* = 25, which is the extinction threshold used throughout the other simulations. Simulations were conducted over 20,000 replicate runs with *K* = 4,000 and a founder population size, *N*_founder_= 500. Recurrent mutation occurs at a rate of *μ* = 0.0000125 per generation. Founder populations originated under conditions of balancing selection via spatial storage effect (light green/dark green, *h*_model_*/h*_drift_ = 1.90 for *P* = 12, *h*_model_*/h*_drift_ = 3.02 for *P* = 24, *h*_model_*/h*_drift_ = 4.78 for *P* = 48), directional selection (pink/purple, *h*_model_*/h*_drift_ = 0.51), or drift (light grey/dark grey). The novel environment was either stable (light green/pink/light grey) with directional selection (*s_d_* = -0.55, *s_b_* = -0.15 with *s*_max_ = 0.0) or perturbed (dark green/purple/dark grey) with temporally varying selection (*s_d_* = -0.55, *s_b_* = -0.15 with *s*_max_ = 0.2).

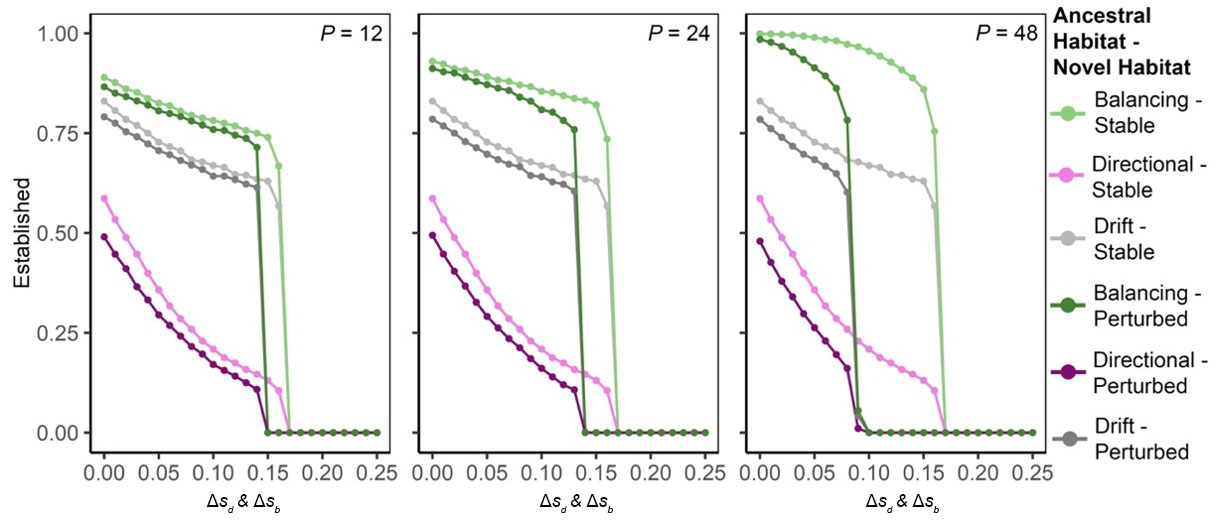

**Figure S3. The probability of establishment in a novel habitat under increased environmental harshness for both alleles.** Simulations were conducted over 20,000 replicate runs with *K* = 4,000 and a founder population size, *N*_founder_= 500. Recurrent mutation occurs at a rate of *μ* = 0.0000125 per generation. Founder populations originated under conditions of balancing selection under the spatial storage effect (light green/dark green, *h*_model_*/h*_drift_ = 1.90 for *P* = 12, *h*_model_*/h*_drift_ = 3.02 for *P* = 24, *h*_model_*/h*_drift_ = 4.78 for *P* = 48), directional selection (pink/purple, *h*_model_*/h*_drift_ = 0.51), or drift (light grey/dark grey). The novel environment was either stable (light green/pink/light grey; *s_d_* = -0.55, *s_b_* = -0.15 with *s*_max_ = 0.0) or perturbed (dark green/purple/dark grey; *s_d_* = -0.55, *s_b_* = -0.15 with *s*_max_ = 0.2). To test different levels of environmental harshness, *s_d_* and *s_b_* were varied simultaneously in increments of −0.01, from −0.55 to −0.80 and from −0.15 to −0.40, respectively (left to right).

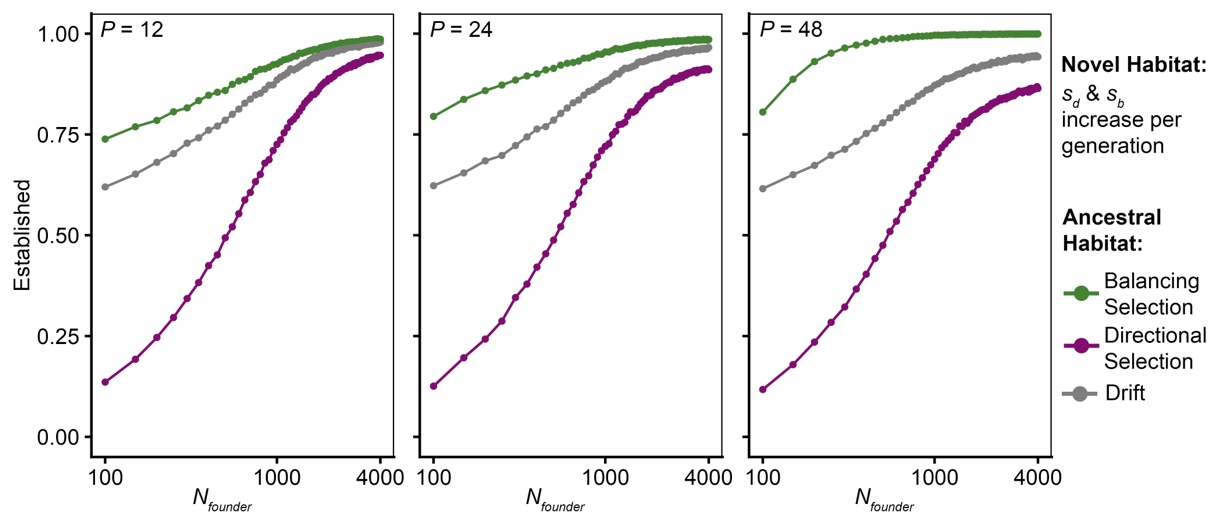

**Figure S4. The probability of establishment in a perturbed novel habitat with an increasing magnitude of varying selection over time as a function of the size of the founder populations, *N*_founder_.** Founder populations originated under conditions of balancing selection under the spatial storage effect (dark green, *h*_model_*/h*_drift_ = 1.90 for *P* = 12, *h*_model_*/h*_drift_ = 3.02 for *P* = 24, *h*_model_*/h*_drift_ = 4.78 for *P* = 48), purifying selection (purple, *h*_model_*/h*_drift_ = 0.51), or drift (dark grey), and experienced a novel habitat with temporally varying selection (*s_d_* = -0.55, *s_b_* = -0.15 with *s*_max_ = 0.2) with *s_t_* based on eq. [5]. Simulations were conducted over 20,000 replicate runs with *K* = 4,000 over 400,000 generations. The *N*_founder_ = 100 to 4000 at increments of 50 individuals (left to right). Recurrent mutation occurs at a rate of *μ* = 0.0000125 per generation. The x-axis is shown on a log scale.

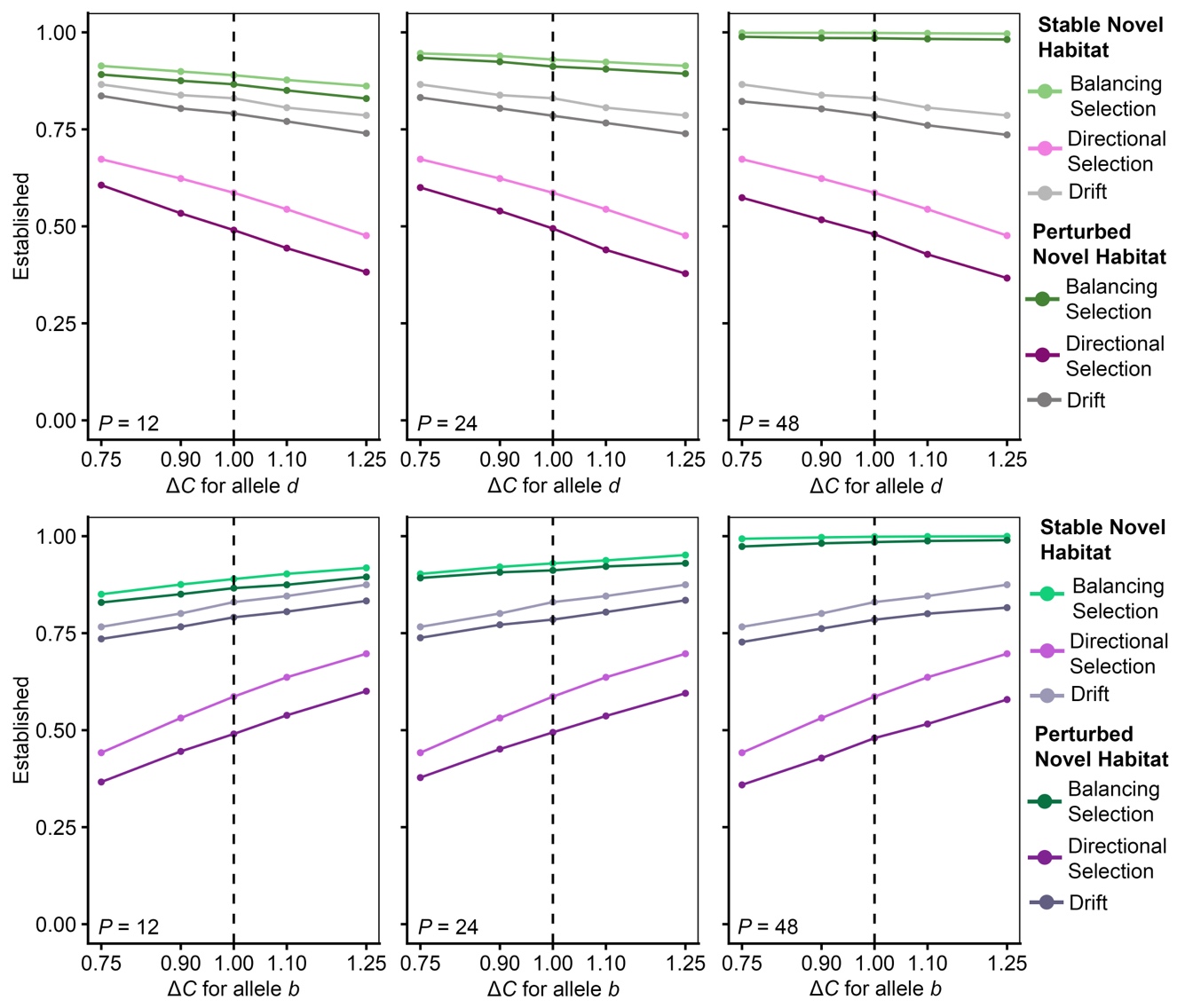

**Figure S5. The probability of establishment in a novel habitat when alleles experience different magnitudes of fitness oscillations.** Simulations were conducted over 20,000 replicate runs with *K* = 4,000 and a founder population size, *N*_founder_= 500. Recurrent mutation occurs at a rate of *μ* = 0.0000125 per generation. Founder populations originated under conditions of balancing selection under the spatial storage effect (light green/dark green, *h*_model_*/h*_drift_ = 1.90 for *P* = 12, *h*_model_*/h*_drift_ = 3.02 for *P* = 24, *h*_model_*/h*_drift_ = 4.78 for *P* = 48), directional selection (pink/purple, *h*_model_*/h*_drift_ = 0.51), or drift (light grey/dark grey). The novel environment was either stable (light green/pink/light grey) with directional selection (*s_d_* = -0.55, *s_b_* = -0.15 with *s*_max_ = 0.0) or perturbed (dark green/purple/dark grey) with temporally varying selection (*s_d_* = -0.55, *s_b_* = -0.15 with *s*_max_ = 0.2). To test unequal magnitude of allele fitness, *s_d_* = -0.55 $\times$*C_d_* or *s_b_* = -0.15 $\times$ *C_b_,* where *C_i_* = 0.75, 0.9, 1.0, 1.1, and 1.25 and *i* = *d*, or *b*, with the other held at *s_t_*.

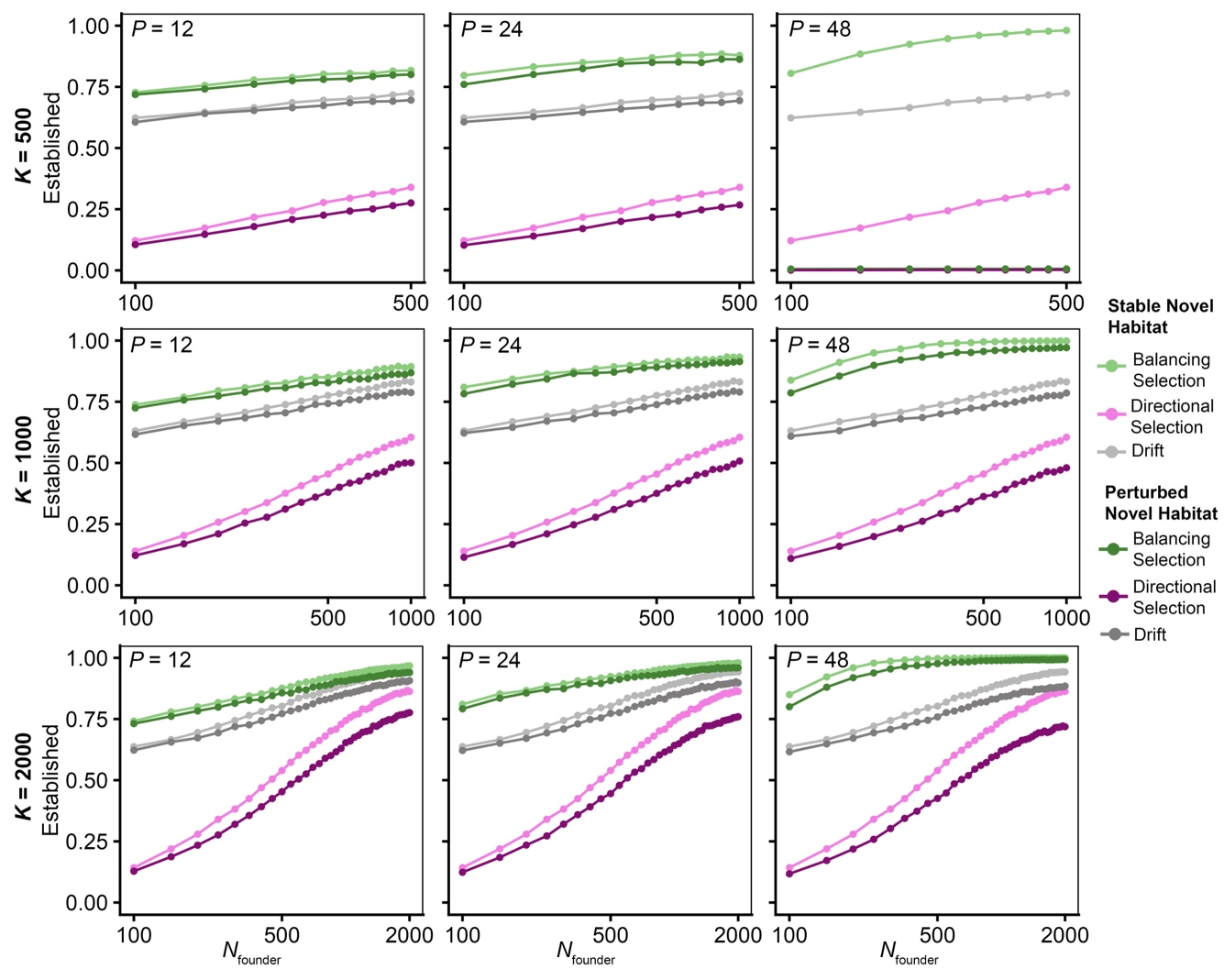

**Figure S6**. **The probability of establishment in a novel habitat under different carrying capacities, *K*** = 500, 1000, and 2000 individuals. Simulations were conducted over 20,000 replicate runs. The *N*_founder_ = 100 to 2000 (or 500, and 1000 when *K* < 2000) at increments of 50 individuals (left to right). Recurrent mutation occurs at a rate of *μK* = 0.05 per generation. Founder populations originated under conditions of balancing selection under the spatial storage effect (light green/dark green, *h*_model_*/h*_drift_ = 1.90 for *P* = 12, *h*_model_*/h*_drift_ = 3.02 for *P* = 24, *h*_model_*/h*_drift_ = 4.78 for *P* = 48), purifying selection (pink/purple, *h*_model_*/h*_drift_ = 0.51), or drift (light grey/dark grey). The novel environment was either stable (light green/pink/light grey; *s_d_* = -0.55, *s_b_* = -0.15 with *s*_max_ = 0.0) or perturbed (dark green/purple/dark grey; *s_d_* = -0.55, *s_b_* = -0.15 with *s*_max_ = 0.2). The x-axis is shown on a log scale.

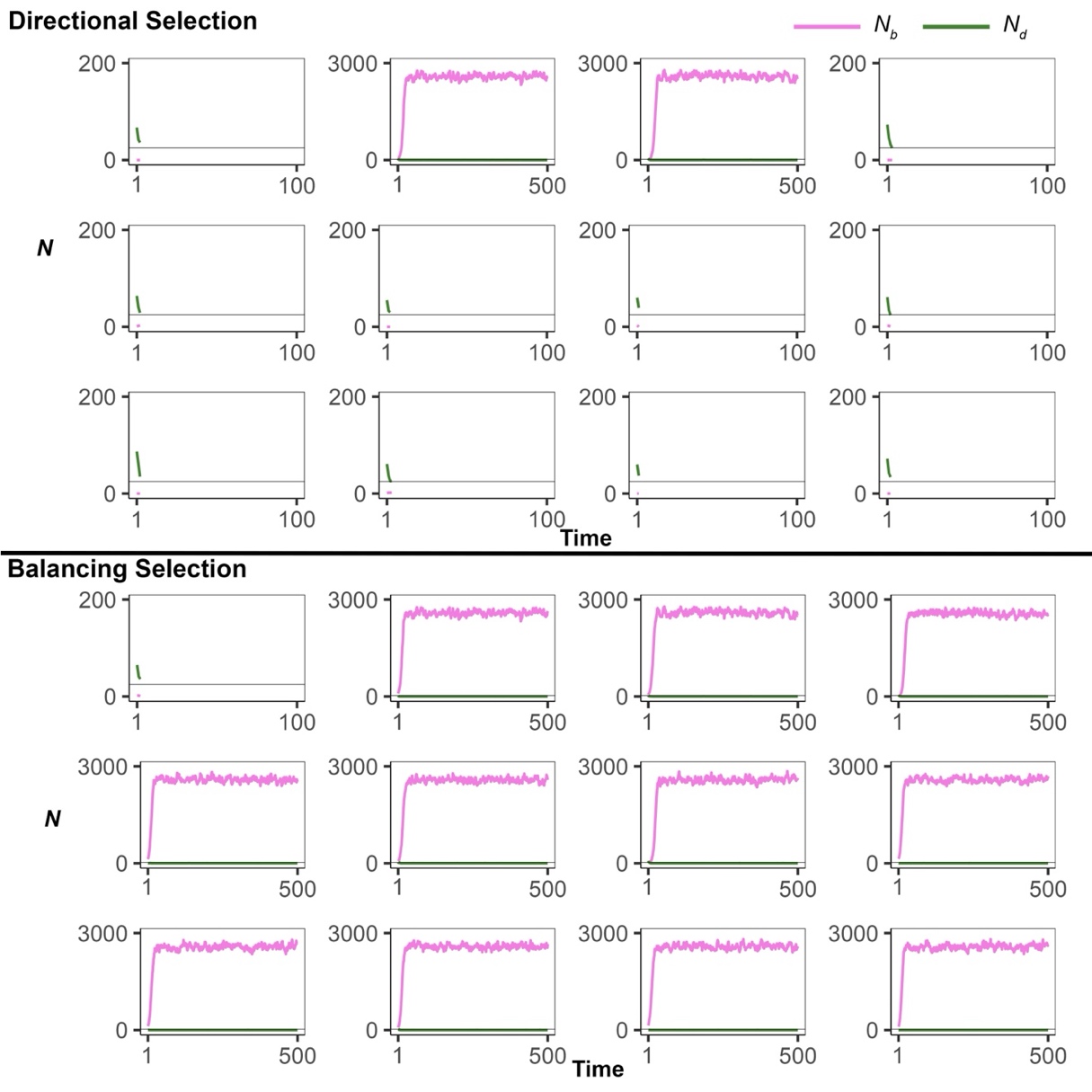

**Figure S7**. **The number of individuals carrying *d* and *b* allele following immigration to a stable new habitat from ancestral populations under purifying selection (top, *h*_model_*/h*_drift_ = 0.51) or balancing selection (bottom, *h*_model_*/h*_drift_ = 4.78)**. Simulations were conducted over 12 replicate runs, with each panel representing a single run, assuming *K* = 4,000 and a founder population size, *N*_founder_ = 100, with *s_d_* = -0.55, *s_b_* = -0.15. Recurrent mutation occurs at a rate of *μ* = 0.0000125 per generation. The number of individuals with the deleterious allele (*N_d_,* shown in green) and the number of individuals with the beneficial allele (*N_b_*, shown in pink) were recorded over 100 or 500 generations, depending on the survival rate. Note that the populations that survived to 500 generations had an increased population size (x-axis maximum increased to 5,000 compared to 200).
